# Dissecting the genetic architecture of quantitative traits using genome-wide identity-by-descent sharing

**DOI:** 10.1101/2021.03.01.432833

**Authors:** Antoine Fraimout, Frédéric Guillaume, Zitong Li, Mikko J. Sillanpää, Pasi Rastas, Juha Merilä

## Abstract

Additive and dominance genetic variances underlying the expression of quantitative traits are important quantities for predicting short-term responses to selection, but they are notoriously challenging to estimate in most non-model wild populations. Specifically, large-sized or panmictic populations may be characterized by low variance in genetic relatedness among individuals which in turn, can prevent accurate estimation of quantitative genetic parameters. We used estimates of genome-wide identity-by-descent (IBD) sharing from autosomal SNP loci to estimate quantitative genetic parameters for ecologically important traits in nine-spined sticklebacks (*Pungitius pungitius*) from a large, outbred population. Using empirical and simulated datasets, with varying sample sizes and pedigree complexity, we assessed the performance of different crossing schemes in estimating additive genetic variance and heritability for all traits. We found that low variance in relatedness characteristic of wild outbred populations with high migration rate can impair the estimation of quantitative genetic parameters and bias heritability estimates downwards. On the other hand, the use of a half-sib/full-sib design allowed precise estimation of genetic variance components, and revealed significant additive variance and heritability for all measured traits, with negligible dominance contributions. Genome-partitioning and QTL mapping analyses revealed that most traits had a polygenic basis and were controlled by genes at multiple chromosomes. Furthermore, different QTL contributed to variation in the same traits in different populations suggesting heterogenous underpinnings of parallel evolution at the phenotypic level. Our results provide important guidelines for future studies aimed at estimating adaptive potential in the wild, particularly for those conducted in outbred large-sized populations.

## Introduction

Given the novel selection pressures stemming from global environmental change, our ability to predict short-term adaptive responses in ecologically important traits in natural populations is becoming ever more important (Waldvogel *et al*. 2020). Through the estimation of quantitative genetic parameters underlying phenotypic traits (*e.g.,* additive genetic variance *V_A_* and narrow-sense heritability *h²*, Lynch & Walsh 1998) a population’s ability to respond to selection can be estimated using various metrics and equations based on these parameters (*e.g*., evolvability, Hansen & Pelabon 2021; multivariate breeder’s equation, Lande 1979; Robertson-Price identity, Robertson 1966, Price 1970). Therefore, estimation of quantitative genetic parameters in the wild is an important, but at the same time, difficult challenge given the logistic demands associated with obtaining the required data.

Traditionally, estimation of *V_A_* and *h²* requires measures of continuous traits among individuals of known relatedness, using specific crossing designs in the laboratory, or pedigree information gathered from natural populations (Lynch & Walsh, 1998). In both cases, establishing the degree of relatedness between individuals is logistically challenging because most non-model organisms are not amenable to laboratory rearing or because building sufficiently large pedigrees requires extensive, and often long-term, work in the wild. However, the genomic era has provided researchers with an increasing number of molecular markers allowing estimation of the actual proportion of genome that is shared identically-by-descent (IBD) between individuals and by directly building the Genomic Relationship Matrices (GRM) from wild individuals genotyped at many marker loci. The GRM can subsequently be included into quantitative genetic models of variance partitioning (*e.g*., animal models, Kruuk 2004) to evaluate the covariance between phenotypic and genetic resemblance. These advances in quantitative genomics have been well explored in the fields of human and medical genetics (Visscher *et al*., 2006, 2007; Manolio *et al.,*, 2009; Visscher, 2009; Powell *et al.,*, 2010; Speed & Balding, 2015), animal and plant breeding (*e.g.* Edwards & Batley, 2010; Hayes & Goddard 2010; Sinclair-Waters *et al.,* 2020), and more recently also in evolutionary ecology (*e.g.*, Sillanpää, 2011; Thompson, 2013; Robinson *et al*. 2013; Perrier *et al.,* 2018; Gienapp *et al.,* 2019; Gervais *et al*. 2020; Duntsch *et al*. 2020; De la Cruz *et al*. 2020; Fraimout *et al*. 2022a). This ‘pedigree-free’ approach has proven particularly useful in getting accurate estimates of additive genetic variance, heritability and genetic correlations underlying the expression of quantitative traits in several wild (or semi-wild) populations including the Soay sheep (Bérénos *et al*. 2014, 2015), Roe deer (Gervais *et al*. 2019) and passerine birds (Robinson *et al*. 2013, Santure *et al*. 2013, Perrier *et al*. 2018).

Nonetheless, and despite having greatly advanced the field of evolutionary genetics, IBD-based approaches to quantitative genetics in the wild carry drawbacks inherent to the need to sample wild individuals. They might also not be suitable for all study systems. The main limitation is that random sampling of wild individuals from populations with high levels of genetic variation increases the probability of sampling unrelated (*i.e*., genetically distant) individuals. If most sampled individuals are unrelated, the statistical power of GRM-based models will be low and heritabilities of focal traits can become underestimated (Ritland 1996, Ritland 2000, Ødergård & Meuwissen 2012, Jensen *et al*. 2014, Gienapp *et al*. 2017). Although this might not be of concern for some moderately closed or philopatric mammalian or avian study systems, or when deep pedigree information is available, studies of species characterized by high fecundity, large effective population sizes or high levels of gene flow could be impaired by the sampling of mostly unrelated individuals.

Here, we explore the utility of IBD-based quantitative genetic models in estimating genetic variance components from large-sized populations with low relatedness among individuals. We hypothesize that the level of relatedness in the population sample and its variance will determine the accuracy of the genetic variance component estimates and that they will be biased downwards if level of relatedness in a population is low. To this end, we estimated quantitative genetic parameters underlying phenotypic variation in a marine outbred population of the nine-spined stickleback (*Pungitius pungitius*). We used empirical data on traits known to be under selection in *P. pungitius* (Karhunen *et al*. 2014), and calculated heritability, additive and dominance genetic variances in controlled crosses of laboratory-raised *P. pungitius* and compared these estimates to those obtained from wild collected samples. Furthermore, we used computer simulations to verify the robustness of our results and further test for the effect of migration rate on the estimation of quantitative genetics parameters. Following our hypothesis, we expect that: i) data from controlled crosses should provide better estimates than wild-collected data and that ii) the accuracy of quantitative genetics parameters should scale negatively with migration rate among populations. In addition, by estimating the contribution of different chromosomes and quantitative trait loci (QTL) to the phenotypic variance of all traits (Visscher *et al*., 2007; Yang *et al.,* 2011) we investigated the genetic architecture of the focal traits. We did this to look for further evidence (cf. Kemppainen *et al*. 2021, Fang *et al*. 2021) that phenotypic parallelism across different *P. pungitius* populations is underlined by lack of similar parallelism in genetic underpinnings of these phenotypic traits.

Our findings suggest that quantitative genomic approaches in the wild should be carried with caution in species where relatedness mean and variance among individuals is expected to be low, that is, in species exhibiting low degree of population structuring and/or high rates of migration.

## Material and methods

### Empirical datasets

Our first objective was to use different data structures and statistical approaches (and see, *Quantitative genetics* section below) to assess their efficiency in obtaining estimates of variance components and heritability of quantitative traits. To this end, we used three different datasets corresponding to three pedigree structures; i) a population sample of wild-collected nine-spined sticklebacks from the Baltic Sea in Helsinki (60°13′N, 25°11′E) brought to the University of Helsinki aquarium facility. This “Helsinki-wild” dataset corresponds to a standard quantitative genomic approach of building GRM from wild-collected individuals of unknown relatedness. ii) Individuals from the Helsinki-wild dataset were used as founders to produce F_1_ generation offspring following standard *in vitro* fertilization procedure for sticklebacks (Divino and Shultz, 2014). F_1_ generation Full-sib/Half-sib families were generated by mating one female to two different males (*i.e*., maternal half-sib design). Once the eggs hatched and larvae started feeding, the families were thinned to approximately 25 offspring per family and moved to two large aquaria so that half of each family were placed in each aquaria unit. The larvae were mass-reared in these aquaria, and their family identity was later identified from the genotype data (see below). After eleven months (May 2013 to April 2014), the experiment was terminated and F_1_ offspring distributed across 87 families were used. This dataset (“Helsinki-HS”) constitutes a classical common garden Full-sib/Half-sib breeding design maximizing the number of different families and relatedness variation in the sample. iii) The third dataset corresponded to a subset of the Helsinki-HS dataset retaining only 46 F_1_ full-sib families (“Helsinki-FS” dataset). The rationale behind generating this dataset was to investigate the impact of half-sib relationships on the estimated parameters (see also *Simulation Study* below).

### Empirical data: Phenotypes

We obtained phenotypic data for all individuals of the three datasets on three morphological traits: body length, body depth and length of the pelvic spines, all known to have heritable basis (Shimada *et al*., 2011; Kemppainen *et al*., 2020) and be involved in adaptive differentiation between freshwater and marine populations (Karhunen *et al*., 2014). Individuals were photographed next to a millimeter scale with a digital camera, and images were used to measure body length and body depth using the tpsDig software (v.2.10; Rohlf, 2006). Body length was measured as the distance between the tip of the snout and the base of the posterior end of the hypural plate. Body depth corresponded to the distance between landmarks 3 and 12 in Herczeg *et al*. (2010). Length of the pelvic spines were measured using digital calipers to the nearest 0.01 mm. We did not focus on the asymmetry of pelvic spines (but see: Blouw & Boyd, 1992; Bell *et al*., 2007; Coyle *et al*., 2007) but rather decided to use the mean values of the left-and right-side measurements as the focal trait. All measurements were made by the same person twice, and the repeatabilities were > 0.9 (*p <* 0.001) for all traits (see also Yang *et al.,* 2016, Kemppainen *et al.,* 2021). Sex information for all samples was obtained first by genotyping all individuals at a sex-specific microsatellite locus (Stn 19; Shikano *et al*., 2011), and subsequently confirmed with sex-specific SNPs. After correcting for missing values, the final datasets contained full phenotypic record for 133 wild individuals (Helsinki-wild, body length only), 936 F_1_ individuals (Helsinki-HS) and 482 F_1_ individuals (Helsinki-FS).

### Empirical data: SNP genotyping

SNP data was the same as in Kivikoski *et al*. (2021). In summary, 49 511 SNPs were obtained from sequencing/genotyping by Diversity Arrays Technology (Pty Ltd, Australia), using their DarTseq technology. Prior to all analyses, we pruned the datasets from sex-linked markers and markers with a minimum allele frequency (MAF) lower than 0.01 using the *raw.data* function of the *snpReady* R package (Granato *et al*. 2018) and included only markers with a maximum of 10% missing data. We further used the *imput* option of the *raw.data* function to allow for data imputation of missing genotypes by using the mean genotypic value at each SNP (Granato *et al*. 2018). This resulted in a total of 15 147 SNPs used for the construction of the GRM (see below).

### Quantitative genetics: Construction of the Genomic Relationship Matrices

We estimated two types of matrices describing the relationship among individuals in each empirical dataset: i) pedigree-based relationship matrices (PRM), corresponding to the theoretical expectation of sibling relatedness based on their pedigree and ii) SNP-based GRM corresponding to the estimated fraction of the genome shared IBD between siblings. The PRMs were constructed from the pedigree structure describing each dataset using the *nadiv* R package (Wolak, 2012) which allowed us to compute both additive (PRM_ADD_) and dominance (PRM_DOM_) matrices using the *makeA()* and *makeD()* functions, respectively.

For the construction of the GRM, we used a modified construct of the additive GRM originally proposed by VanRaden (2008):

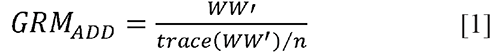

where *W* is the marker matrix of additive coefficients and *n* the number of individuals, as implemented in the *snpReady* R package (Granato *et al*., 2018) for each dataset to construct GRMs from autosomal loci. This allowed us to produce for each dataset both additive and dominance variance matrices (GRM_ADD_ & GRM_DOM_; Granato *et al*., 2018). The construction of GRM_DOM_ follows the same formula as in equation [1] replacing the matrix *W* by the matrix *S* corresponding to the dominance deviation coefficients (see p. 5 in Granato *et al*. 2018).

### Quantitative genetics: Estimation of variance components and heritabilities

Three quantitative genetic parameters underlying trait variability were estimated: additive genetic variance (V_A_), dominance variance (*V_D_*) and heritability (*h^2^*). We employed the ‘animal model’ approach (Kruuk, 2004), a random effect model using the PRM or GRM as the covariance of the random effect linking to the phenotypes, to estimate these parameters using the relatedness among individuals in each dataset characterized in each relationship matrix. The Bayesian approach implemented in the *MCMCglmm* R package (Hadfield, 2010) was used to fit the following model for each trait and each dataset separately:

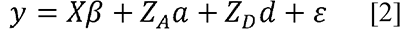

where *y* is the vector of phenotypic values for each trait, *β* is the vector of fixed effects, *a* and *d* are the vectors of random additive and dominance effects, respectively. ε is the vector of residual errors and *X*, *Z_A_* and *Z_D_* the design matrices relating to the fixed effects and the additive and dominance random effects, respectively. We added the sex of the individuals as a fixed effect and implemented additive and dominance effect from either type of relationship matrix (PRM or GRM) as random effects. Model convergence was checked visually from the mixing of MCMC chains by inspecting trace plots of many different model parameters, using the Heidelberger and Welch convergence test (Plummer *et al*. 2006). We assessed whether models had been run for enough iterations by inspecting the effective sample sizes of the MCMC posterior distributions of each variance components. We calculated heritability as the ratio of *V_A_* to total phenotypic variance (*V_P_*) for a given trait. For each model, we evaluated the uncertainty in the estimation of variances and heritability based on the 95% Highest Posterior Density (HPD) intervals constructed from the posterior distribution of each variance component using the *HPDinterval()* function (Plummer *et al*. 2006). We then compared the 95% HPD intervals for all parameters (*V_A_, V_D_*, *h²*) between models to assess the precision of each approach in estimating quantitative genetic parameters. As the pedigree for the Helsinki-wild data contains no information (*i.e.,* all individuals are considered unrelated) we only ran models based on the GRM for this dataset. All variance components obtained with *MCMCglmm* using GRMs and PRMs are reported as the median of the posterior distributions for each model.

Finally, meaningful comparisons of variance components estimated using different relationship matrices (here, GRM & PRM) must refer to a common reference population wherein genetic variance is estimated (Speed & Balding 2015, Legarra 2016). Specifically, models using PRM will estimate variance in the base population of the pedigree (*i.e.,* the unrelated founders) while models using GRM will do so within the set of genotyped individuals (Powell *et al*. 2010, Legarra 2016; Joshi *et al*. 2020).

Here, we followed the approach of Legarra (2016) and adjusted all variance components as follows:

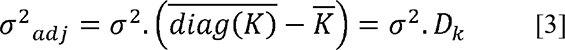

where σ² is the variance component estimated from the animal model and *K* the corresponding relationship matrix (Legarra 2016; Joshi *et al.,* 2020). By doing so, we set a reference population to provide meaningful comparison between the PRM and GRM-based models.

### Genetic architecture: Chromosome partitioning of the genetic variance and QTL mapping

To investigate the genetic architecture of the three quantitative traits, we estimated the proportion of phenotypic variance explained by SNPs from each separate chromosome. To this end, we used the Genomic Best Linear Unbiased Prediction (G-BLUP) approach from the GCTA software (Yang *et al*. 2011). We calculated a GRM for each chromosome and a GRM for all but the focal chromosome and estimated genetic variance parameters with the GREML approach implemented in GCTA. Models were run independently for each trait and each chromosome for all crosses.

Furthermore, we performed QTL mapping using three inter-cross F_2_ datasets (see *Supplementary methods*). We used a single-locus mapping approach (Li *et al*. 2017, 2018) to identify the QTL explaining variation in body size, body depth and pelvic spine length. The total phenotypic effect of each SNP can be obtained from the regression coefficient of the Gaussian regression model:

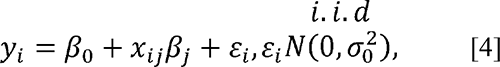

where *y_i_* is the vector of phenotypes for individuals *i*; *β_0_* is the phenotypic mean*; x_ij_* is the genotypic value of individual *i* and marker *j* coded as −1, 0 and 1 for the three genotypes AA, AB and BB, respectively; *β_j_* is the a normal distribution with zero mean and unknown variance 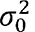. In our inter-cross F_2_ design, up to four possible effect of the SNP *j*, and ε_i_ is the residual error assumed to be independent and identically distributed (*i.i.d.*) under segregating alleles can be found: two alleles A1 and A2 from the dam, and two alleles B1 and B2 from the sire (Xu, 1996, 2013). Thus, model [4] can be reformulated as:

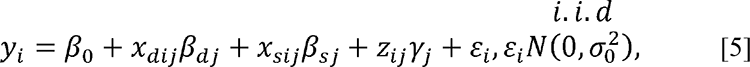

where the substitution effects *β_dj_* and *β_sj_* correspond to the alleles A1 and A2 and B1 and B2 at locus *j* for the dam *d* and sire *s*, respectively, and where *γ_j_* is the dominance effect. In model [5], the genotype matrix coding system for *x_dij_*, *x_sij_* and *z_ij_* (Xu, 2013) is specified as:

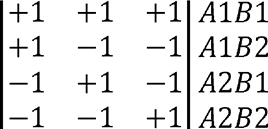

To obtain additional information on the origin of the QTL effects and distinguish between the dam and sire allele effects, parental phasing information needs to be incorporated in the model [5]. To this end, we used the data produced in Kemppainen *et al*. (2021). Briefly, parental and grandparental phases were obtained from a dense SNP panel (see detailed description in Kemppainen *et al*. 2021 and Li *et al*. 2018) using the LEP-MAP3 software (Rastas, 2017). Data redundancy due to linkage was reduced using a linkage disequilibrium (LD) network approach implemented in the *LDna* R package (Kemppainen *et al*. 2015). Each LD-cluster comprising set of highly correlated SNPs was subjected to a principal component analysis (PCA) and the PC-coordinates from the first axis explaining the largest proportion of variation was used for QTL mapping. For each of the three F_2_ crosses, we applied the model from equation [5] separately to each phenotypic trait using the complexity-reduced SNP panel and sex as covariate. A permutation procedure (10,000 permutations) was used in each QTL model to control for false positives (Li et al., 2017; Westfall & Young, 1993).

### Simulation study

We ran forward-in-time, Wright-Fisher simulations with Nemo (v.2.3.56; Guillaume & Rougemont 2006) to generate genomic and phenotypic data for datasets varying in their pedigree structures. From a base population of *N* = 10,000 individuals, three categories of datasets matching our empirical data were generated after 10N generations: i) wild samples, ii) full-sib/half-sib crosses and iii) full-sib crosses. For each cross type we generated different sample sizes to evaluate the number of individuals needed to obtain precise variance component estimates. For the full-sib crosses, we simulated a dataset with three independent (i.e., unrelated founders) full-sib families each consisting of 200 F_1_ offspring (“FS_3×200”, *N* = 600) and a dataset with five independent full-sib families each with 200 F_1_ offspring (“FS_5×200”, *N* = 1000) and finally, a dataset equivalent to our empirical dataset consisting of 50 families each containing 20 offspring (“FS_50×20”, *N* = 1000). For the full-sib/half-sib design we simulated a design consisting of 50 dams each mated to two sires and each family consisting of either five F_1_ offspring per sire (“HS_50×10”, *N* = 500) or 10 F_1_ offspring per sire (“HS_50×20”, *N* = 1000) and an additional design with 150 dams each mated to two sires and each family consisting of three F_1_ offspring (“HS_150×6”). Finally, we simulated two datasets of wild unrelated individuals consisting of 500 (“Wild_500”) or 1000 individuals (“Wild_1000”) randomly sampled from the base population.

Moreover, we explored the effect of variation in relatedness on the estimation of variance components by simulating additional ‘wild’ datasets with varying levels of IBD among individuals. To this end, we simulated a structured population (*N* = 10,000) of 5 and 10 sub-populations connected by migration in a Stepping-Stone model. Migration rates varied between low (*N_e_m* = 1.6, 3.2) and high (*N_e_m* = 16, 32) (Fig. S1) so that relatedness among all individuals would scale negatively with migration. In other words, under a low-migration regime, individuals within sub-populations are more related to one-another (Fig. S1a, c), while high migration rates homogenize genetic variation in the meta-population and consequently, decreases average relatedness (Fig. S1b, d). After 10N generations of random mating (Wright-Fisher) and migration, we generated samples of 500 individuals equally sampled among sub-populations (100 or 50 individuals per sub-population).

For each dataset, a polygenic quantitative trait underlined by 100 biallelic QTL was simulated and 10,000 neutral biallelic SNP markers were generated on the same genetic map for estimation of genomic relatedness. Loci were placed on 20 chromosomes with total map length 190cM. Neutral markers were placed equidistantly every 0.5cM, while QTL locations were randomly set. QTL allelic effects were ±0.4 with mutation rate 10^-4^. Mutation rate for neutral markers was 10^-5^. Narrow-sense heritability *h^2^* was maintained constant across simulations by adjusting the environmental variance of the trait *V_E_* to the within population additive genetic variance *V_A_* stemming from mutation-drift(-migration) balance. GRM_ADD_ for the simulated genotypes were constructed using the same approach as described above after pruning for markers with a MAF lower than 0.01. We used the same modeling approach to estimate all variance components from each simulated dataset with *MCMCglmm* as described above. For each dataset, 10 replicates were generated and the mean values of all replicates along with their standard deviations are reported.

## Results

### Distribution of IBD-sharing values

The GRM estimated from the Helsinki-wild cross data was well representative of the full-sib/half-sib structure of the data (Fig. S2) with a high frequency of full-sibs sharing half of their genome IBD (0.5; Fig. S2) and half-sibs showing an average relatedness coefficient of 0.25. Most founders were ‘unrelated’ (IBD sharing < 0.1; Fig. S2), but we did find some degree of relatedness among parents despite the random nature of the sampling (Fig. S2).

### Quantitative genetics parameters

Overall, more precise estimates of quantitative genetic parameters were recovered from the half-sib/full-sib data (as represented by the width of the 95% HPD intervals) compared to those obtained using either unrelated individuals from the wild, or pedigree information (Fig. 1, Table S1). In the Helsinki-wild dataset, estimates of quantitative genetic parameters for body size could not be recovered using the GRM, and genetic variance component estimates, as well as heritability estimates, had wide 95% HPD intervals (Fig. 1, Table S1). We found similar moderate heritability for the three quantitative traits (Fig. 1) but different levels of underlying additive genetic variance, with SL showing the highest and PL the lowest V_A_ (Fig. 1). Overall, the contribution of dominance genetic variance (V_D_) in all traits for these datasets was small (Fig. 1). Accuracy of the parameter estimation using the Helsinki-FS data was on par with the Helsinki-HS dataset, but the latter tended to be more precise (i.e., with lower HPD intervals) for all variance component estimates (Fig. 1).

**Figure 1.**
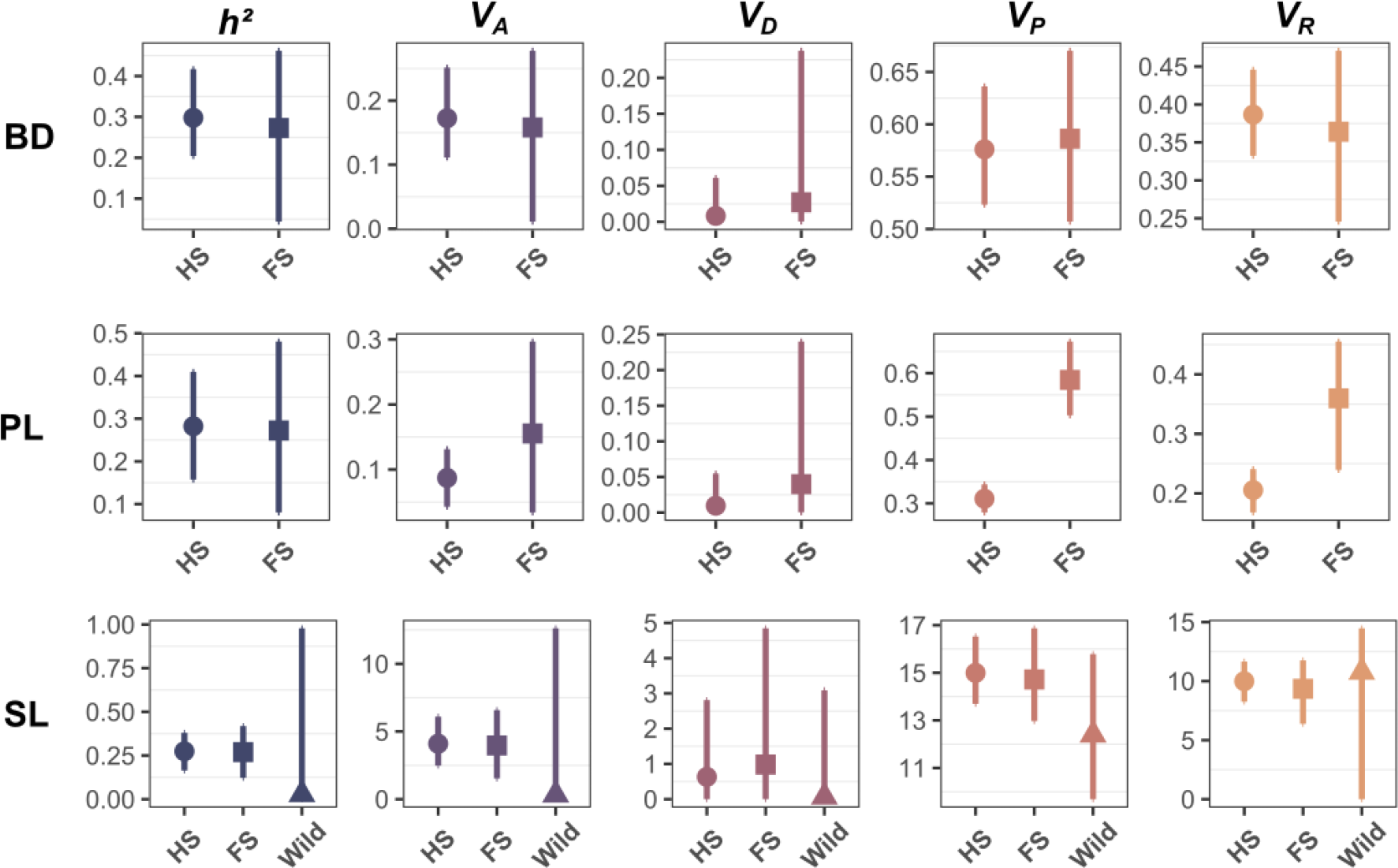
Quantitative genetic parameters estimated from the empirical datasets. The narrow-sense heritability (*h²*), additive genetic variance (*V_A_*), dominance variance (*V_D_*), phenotypic variance (V_P_) and residual variance (*V_R_*) estimated for the empirical datasets are shown. For each dataset and each variance component, the median of the posterior distribution is shown. The 95% HPD interval is represented by vertical solid lines. For each trait (SL: standard length; BD: body depth; PL: pelvic length) each variance component is shown as estimated from the half-sib/full-sib design (HS, filled circle), the full-sib design (FS, filled square) and from the wild population sample (Wild, filled triangle).

Results of the simulation study showed that *V_A_* and *h²* were highly underestimated in all but the half-sib/full-sib crosses (Fig. 2). Accuracy of the estimation (i.e., distance of the point estimates from the true simulated value) increased with increasing family numbers, while precision (i.e., width of the confidence intervals) increased with increasing sample size (Fig. 2). As predicted, and despite the relatively large sample sizes, wild samples did not allow for meaningful estimation of *V_A_* or *h²*.

**Figure 2.**
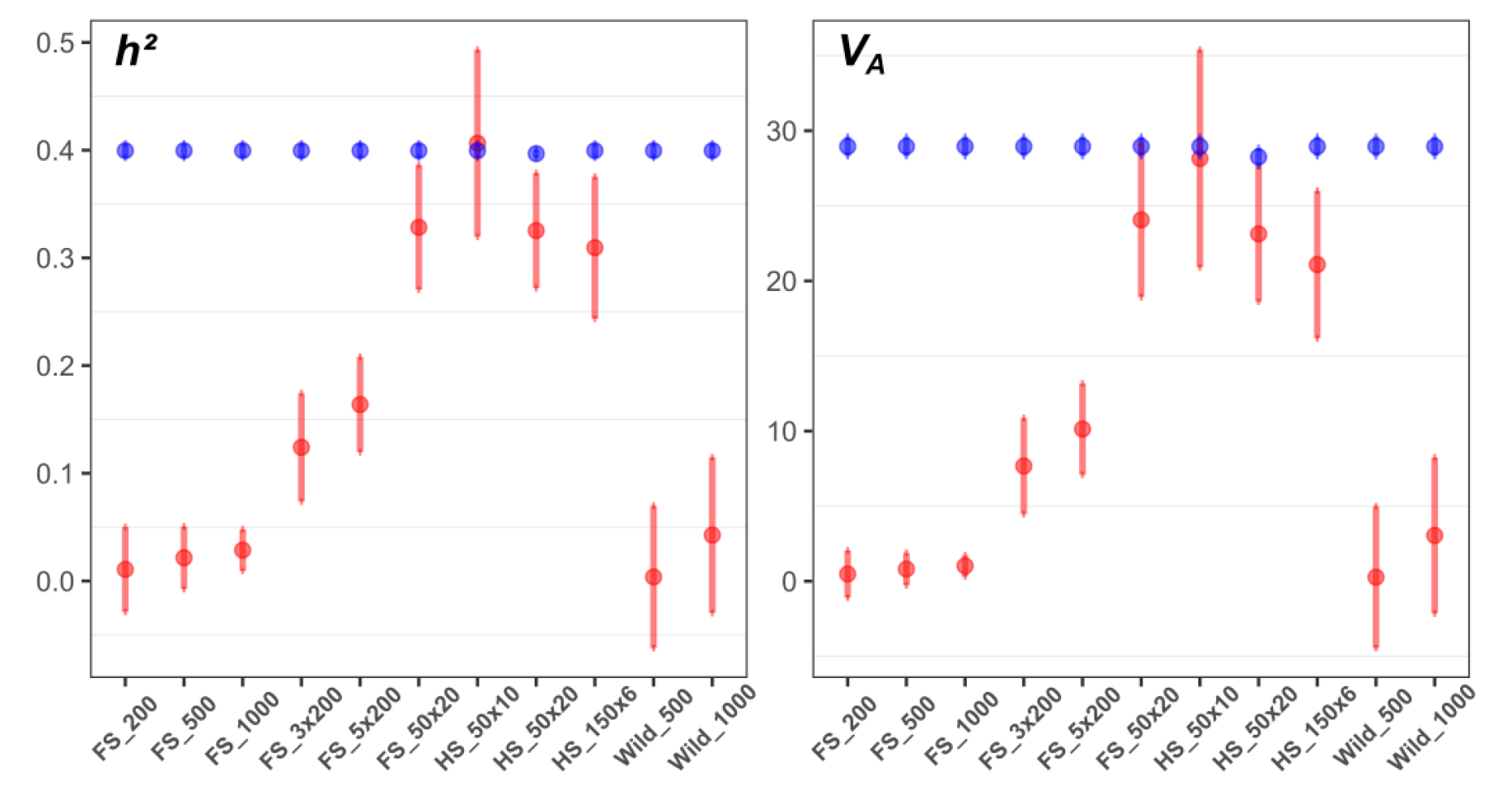
Quantitative genetic parameters estimated from the simulated datasets. The narrow-sense heritability (*h²*) and additive genetic variance (*V_A_*) estimated for the simulated datasets are shown. For each dataset and each variance component, the mean estimate over all replicates is shown (filled circle) along with the 95% HPD interval (vertical bars) for the true simulated values (blue) and the estimates recovered from the *MCMCglmm* models (red). Codes for the datasets on the x-axis are explained in *Methods*.

Further investigations of the wild datasets showed a clear effect of migration rate and relatedness levels on the statistical power to estimate *V_A_* and *h²* (Fig. 3). Migration rates among the subpopulations in each simulated meta-population scaled negatively with the accuracy of *V_A_* and *h²* parameters and the most accurate estimates were obtained using the samples with the lowest migration rate and therefore, the highest level of relatedness (Fig. 3, Fig. S1).

**Figure 3.**
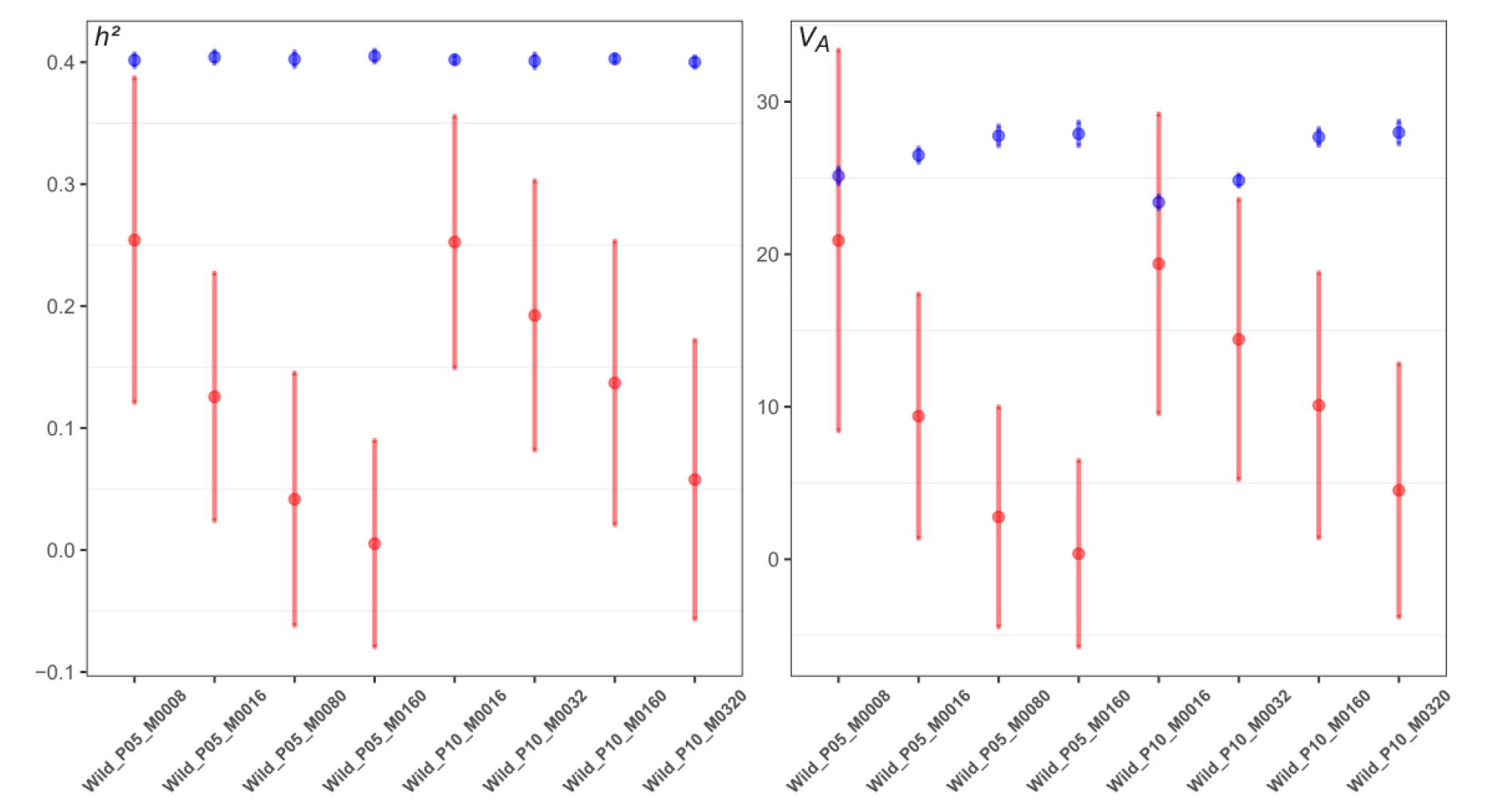
Effect of migration rate on the quantitative genetic parameters estimated from the wild simulated datasets. The narrow-sense heritability (*h^²^*) and additive genetic variance (*V_A_*) estimated for the wild simulated datasets are shown. For each dataset and each variance component, the mean estimate over all replicates is shown (filled circle) along with the standard deviation (vertical bars) for the true simulated values (blue) and the estimates recovered from the *MCMCglmm* models (red). Codes for the datasets on the x-axis correspond to the type of metapopulation used (P05, P10; 5 or 10 subpopulations, see *Methods*) and the level of migration rate among subpopulations (M).

### Genetic architecture: Chromosome partitioning

Overall, the proportion of variance explained by each individual chromosome for each trait was low to moderate in all crosses (Fig. 4, 5), except for pelvic spine length (PL) in the HEL x RYT cross (Fig. 5) where Chromosome 7 explained 66% (SE = 0.086) of the total phenotypic variance. In the Helsinki-HS, we found significant regional heritability for body depth and pelvic length in Chromosomes 6 and 5, respectively (Fig. 4). In the F_2_ crosses, different chromosomes contributed to the total phenotypic variance of all traits in the three datasets (Fig. 5).

**Figure 4.**
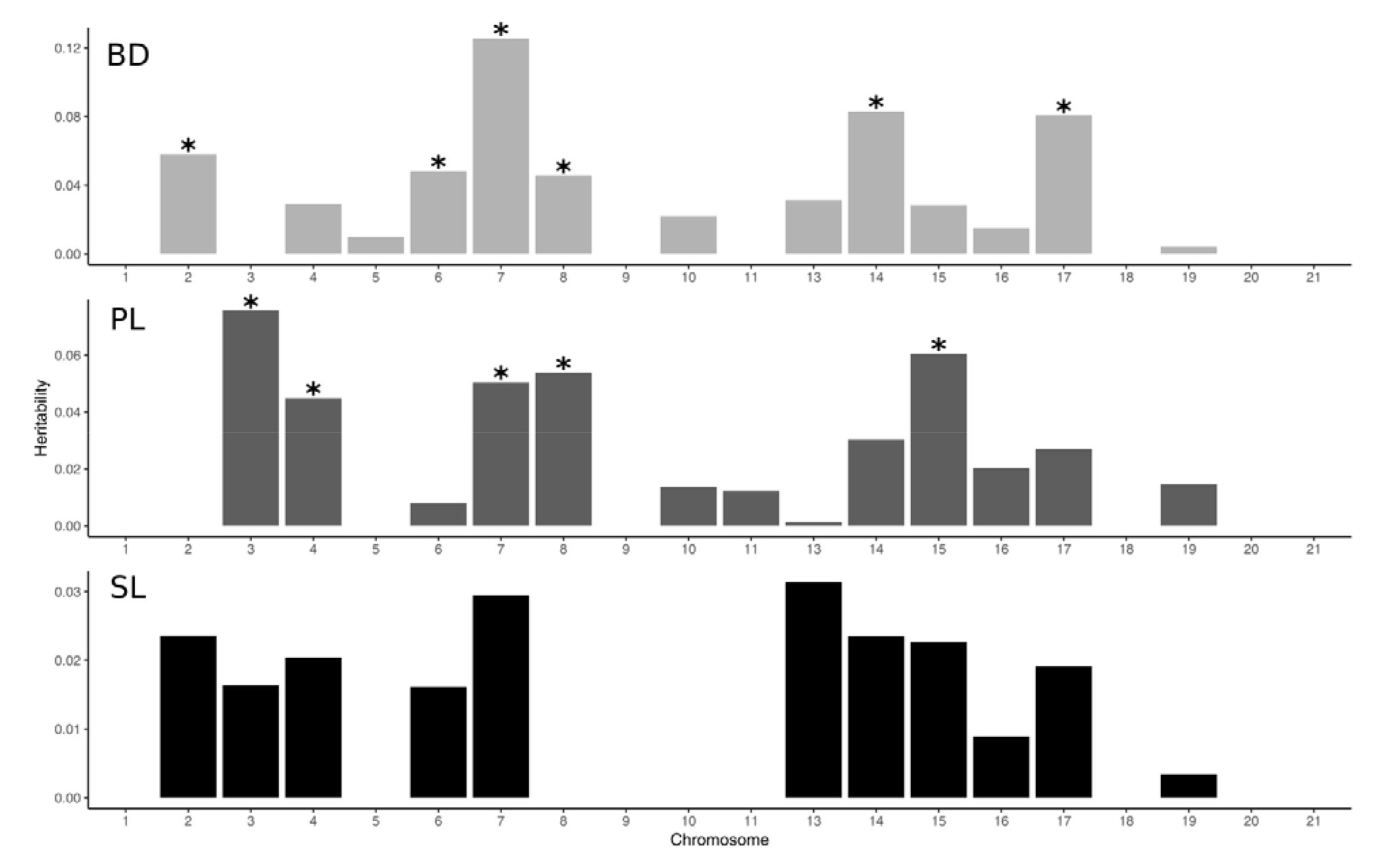
Chromosome partitioning of the genetic variance (Helsinki). The phenotypic variance in body depth (BD), pelvic spine length (PL) and standard length (SL) explained by the SNPs are shown of each chromosome for the Helsinki crosses. Black stars correspond to non-zero estimates (+/-SE) for each chromosome.

**Figure 5.**
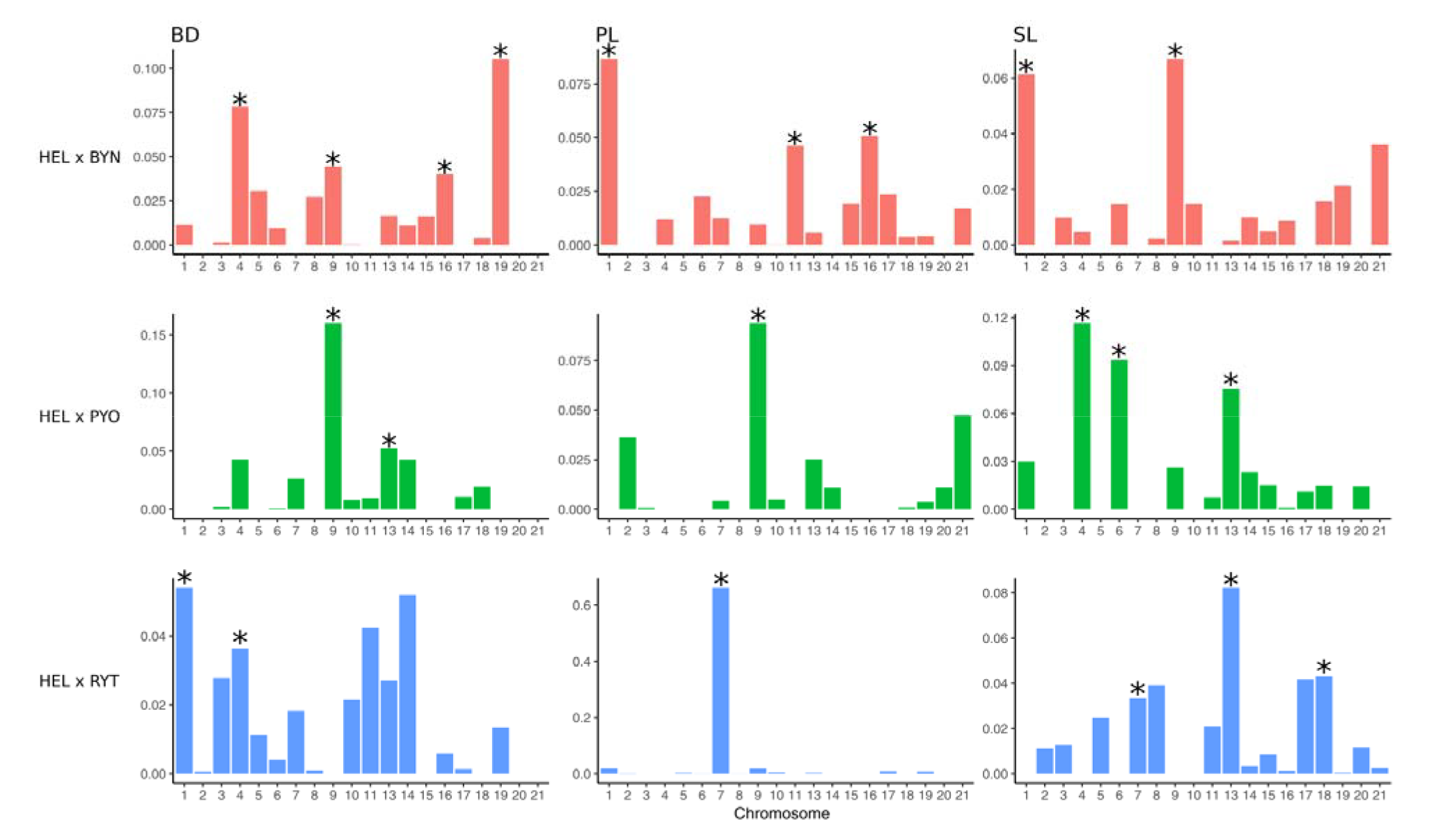
Chromosome partitioning of the genetic variance (QTL crosses). The phenotypic variance in body depth (BD; left column), pelvic spine length (PL; center column) and standard length (SL; right column) explained by the SNPs are shown of each chromosome for the HB cross (red; top row), HP cross (green; middle row) and HR (blue; bottom row). Black stars correspond to non-zero estimates (+/-SE) for each chromosome.

The proportion of variance explained by each chromosome was significantly correlated with chromosome length for body size only in the HEL x PYO cross (*r* = 0.619, *p* = 0.003; see Fig. S3 and Table S2 for details on chromosome length).

### Genetic architecture: QTL mapping

The genetic architecture of the three focal traits differed between the three different F_2_ crosses (Fig. 6). In the HEL x BYN cross, we found no significant QTL underlying the variation in standard length (Fig. 6A), one significant QTL of parental origin (*i.e.,* from the F_0_ pond sire) for body depth on Chromosome 16, and three significant QTL for pelvic spine length on Chromosomes 15, 16 and 21 also of a paternal origin (Fig. 6A). In addition, there was one significant QTL of maternal origin (*i.e*., F_0_ marine female) on Chromosome 6 (Fig. 6A). In the HEL x RYT cross, a single significant QTL underlying the variation in pelvic spine length on Chromosome 7 (Fig. 6B) inherited from both F_0_ parents was found. This QTL expressed also dominance on this trait (Fig. 6B). In the HEL x PYO cross, there was a single paternal QTL for standard length located on Chromosome 1 and one significant QTL for the dominance effect on Chromosome 9. A large effect QTL of shared paternal, maternal and dominance origin located on Chromosome 9 was found to explain variation in body depth (Fig. 6C). Finally, a single significant QTL on Chromosome 9 explained the phenotypic variation in pelvic spine length in this cross (Fig. 6C).

**Figure. 6.**
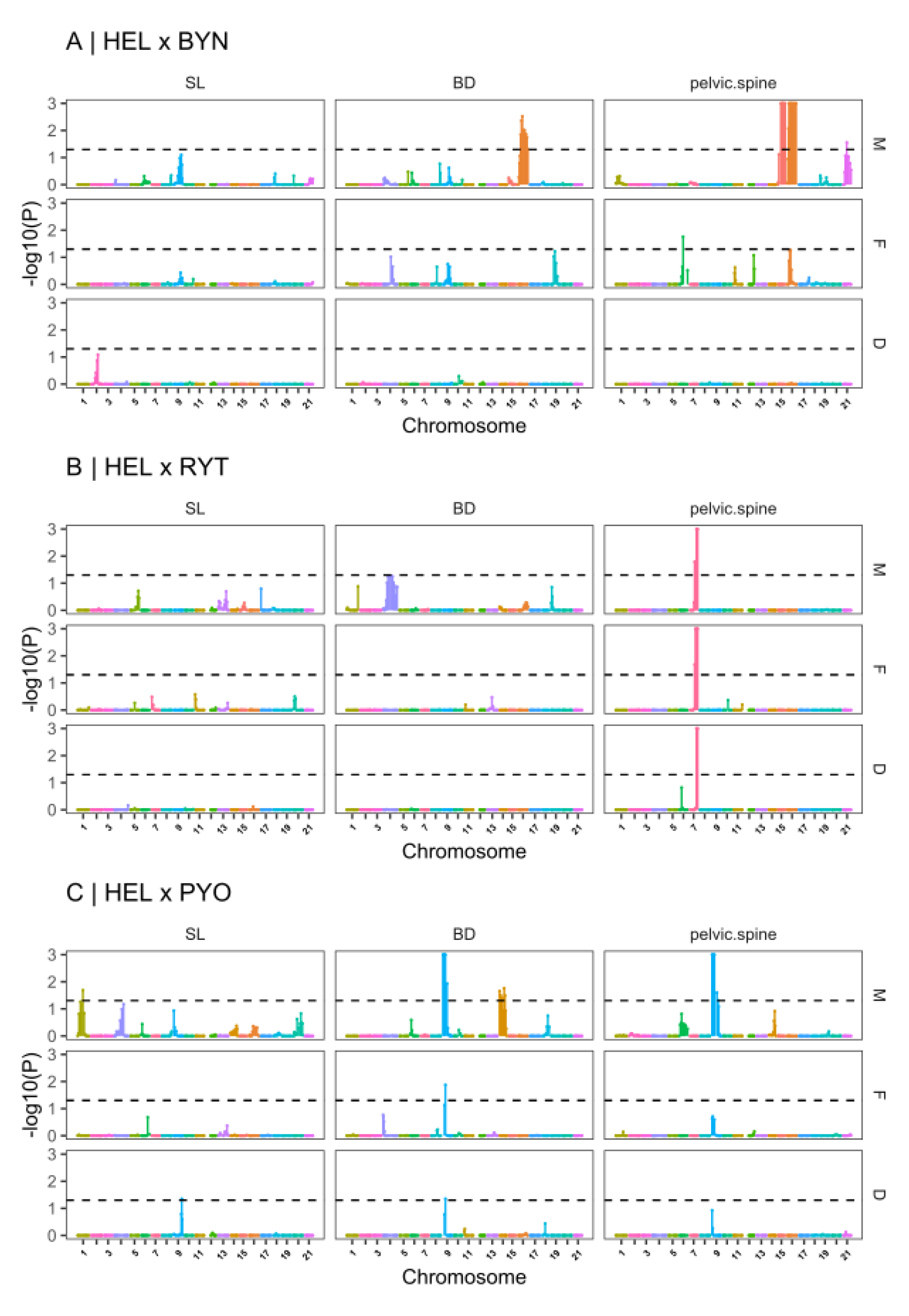
QTL-mapping results for the three phenotypic traits. Results from the four-way QTL-mapping are shown for each cross (A, B, C) and for each trait (SL: standard length; BD: body depth; PL: pelvic spine length). For each cross and trait, panels show whether the QTL is inherited from the sire (M) or the dam (F) along with the dominance effect (D) estimated from model [5] in the main text. Results are based on permutation and the significance threshold (dashed horizontal line) is shown on the logarithm scale (*p* = 0.05). Colors represent different chromosomes.

## Discussion

The results show that genome-wide IBD information derived from SNP markers using controlled crosses such as half-sib/full-sib family can yield meaningful estimates of quantitative genetic parameters for ecologically important traits, and greatly improve the precision of estimates compared to traditional pedigree-based approaches and to estimates obtained from wild unrelated individuals. Moreover, implementation of the GRM in the animal model allowed for greater model complexity, particularly by adding a random effect term for the estimation of dominance genetic variance. In line with the results of some earlier studies, which have estimated dominance variance components for morphological traits (Roff & Emerson 2007, reviewed in Wolak & Keller 2014), the dominance contributions for all traits were negligible in our data. In the following, we discuss the usefulness and limitations of our results, and their potential utility for evolutionary studies of wild populations.

### Quantitative genomics in the wild

Despite the appealing prospect of applying GRM-based animal models directly to samples originating from the wild rather than from complex laboratory crosses, our results show, in line with early predictions (Ritland 1990, Ritland *et al*. 1996), that such approach should not be undertaken in populations with low variance in relatedness. This was particularly evident from our simulation study showing that estimates of heritability and additive genetic variance were severely underestimated when using unrelated individuals with low variance in relatedness compared to half-sib/full-sib data. Furthermore, the results show that in a simulated meta-population representative of a wild outbred species, migration rate (or inversely, population structure) would have to be very low, and variance in relatedness among individuals to be substantial in order to obtain accurate estimates of *V_A_* and *h²*. Of course, the used simulation scheme makes several assumptions (e.g., random mating and absence of mate choice) that are not necessarily met in all species with more complex mating strategies and population structures. Furthermore, sweepstake reproductive success typical of marine species (Vendrami *et al*. 2021) can disrupt the genetic homogeneity of populations otherwise characterized by large census population size, and, therefore, increase the mean and variance in relatedness at local geographic scales. Nonetheless, the simulations were parameterized with results from real genomic data derived from the study of Baltic Sea nine-spined sticklebacks (Fang *et al*. 2021, Feng *et al*. 2022). Hence, our results can be considered representative of at least this outbred fish species, and likely other marine organisms with large population sizes. As such, our results provide important guidelines for future studies aiming at estimating quantitative genetic parameters such as marine species and more generally in species characterized by large population sizes.

### Distribution of IBD values

In humans, relatedness among full-sibs can vary substantially around the 0.5 expectation: Visscher *et al*. (2006) reported IBD sharing among full-sibs to range from 0.317 to 0.617. Here, we found the average IBD sharing to span much wider range than in humans (mean IBD sharing: 0.509, min: 0.259; max: 0.737; Fig. S2). In other words: some full-sib pairs did not share more of their genome than half-sibs, while others shared more than expected from brother-sister mating. Variance in IBD sharing between sibs is inherently linked to structural properties of the genome (i.e., number of chromosomes, chromosome length) and the recombination process and number of crossovers occurring during meiosis: higher number of crossovers will lead to lower variance in IBD (Risch & Lange, 1979; Kivikoski *et al*., 2021). Linkage maps constructed from human data are on average twice the length as that from *P. pungitius* (Broman *et al*., 1998; Kivikoski *et al*., 2023) indicating more crossovers in the human genome than in that of the nine-spined stickleback. Therefore, it is possible that this difference in the recombination landscape between the two species along with the smaller size of *P. pungitius*’ genome (Xu 2006) is responsible for the different range of IBD proportions in sticklebacks. Regardless of the mechanism behind the higher variance in the nine-spined stickleback IBD, our results confirm that this species, and likely also its close relative, the three-spined stickleback (*Gasterosteus aculeatus*), sharing similar genetic architecture and cross-over pattern (Kivikoski *et al.,* 2023), are particularly suitable models for quantitative genomic studies of wild populations (Merilä, 2013). This view is further reinforced by the discovery that the two species differ fundamentally in the way standing genetic variation is distributed among local populations, and therefore in the way local adaptation is expected to proceed in response to similar selection pressures in different populations (Fang *et al*. 2021).

### Dominance variance underlying quantitative traits

The results further demonstrate that use of the GRMs in our models allowed for greater model complexity by implementation of the dominance matrix (GRM_DOM_) as an additional random term to estimate *V_D_*. To date, dominance genetic variance has received less attention in the evolutionary genetic literature (Wolak & Keller, 2014). This is in part due to the fact that founding theoretical work predicted that non-additive genetic effects should contribute little to quantitative trait variance (Fisher, 1958), a prediction later verified in some empirical studies (*e.g.*, Hill *et al.,* 2008; Zhu *et al.,* 2015; Class & Brommer, 2020; but see: Merilä et al. 2004). Nevertheless, a body of theoretical and empirical work suggest that dominance variance can indeed account for a significant proportion of phenotypic variance in the wild when populations are finite and genetically structured (*e.g.*, Wright, 1931; Crnokrak & Roff, 1995; Kosova *et al.,* 2010). Hence, the importance (or lack thereof) of dominance genetic variance cannot be taken for granted and requires empirical data. Two results of particular importance regarding dominance variance from our study are worth highlighting. First, we found that dominance genetic variance accounted for a very low proportion of the total phenotypic variance for all traits in all datasets. On average, it accounted for 0.19% of *V_P_* across datasets compared to *V_A_*, which explained on average 21.2% of *V_P_*. Thus, our results are in line with some earlier studies suggesting that dominance variance in morphological traits can be negligible. Second, the results further demonstrate the difficulty in obtaining estimates of *V_D_* from family data. Here, even when using an appropriate crossing design (*i.e.*, maternal half-sib; Wolak & Keller, 2014) with relatively large number of families and individuals, we found that estimates of *V_D_* obtained from pedigree-based analyses tended to be overestimated as compared to the ones obtained with GRM. As a result, uncertainty around the estimation of the other components of variance in animal model was affected, and heritability estimates became biased downwards. All this said, we acknowledge that there are more advanced breeding strategies combining paternal half-sibs and double-first-cousins that are tailored for dominance variance estimation (Sztepanacz & Blows, 2015; Koch *et al*. 2020) and that larger sample sizes than those used here might be required to obtain reliable estimates of *V_D_*. While it is possible that we did not have enough statistical power to accurately estimate dominance variance in our data, the range of possible values for our focal traits should nonetheless be captured by the HPD intervals of our model estimates.

### The genetic architecture of parallel evolution in P. pungitius

The pace of adaptive evolution is determined by the amount of heritable variation in traits under selection, but the heritability of a given trait may be specific to a given environment, or sex (Price & Schluter, 1991; Roff, 1997; Hoffmann & Merilä, 1999; Wilson *et al*., 2010). Here we studied three morphological traits previously shown to be involved in the phenotypic differentiation between different ecotypes of *P. pungitius*. Following the colonization of lakes and ponds from the marine environment, freshwater populations of *P. pungitius* have repeatedly evolved distinct phenotypes such as increased body size (*i.e.*, gigantism; Herczeg *et al*. 2009), changes in behavior (Herczeg *et al*. 2009, Fraimout *et al*. 2022b) and pelvic reduction (Kemppainen *et al*. 2021). A reasonable assumption is therefore that sufficient additive genetic variance was available in the marine ancestral population to respond to the selection pressures associated with the colonization of the freshwater habitat. In the current study, the Helsinki population may be considered as representative – *i.e.*, genetically and morphologically similar (Shikano *et al.,* 2010, Karhunen *et al.,* 2014) – of an ancestral marine population of *P. pungitius,* and in line with the previous assumption, we found significant *V_A_* underlying all three traits in this population. Further dissection of the genetic architecture of body size revealed that heritability of this trait was partitioned across the genome, with several chromosomes accounting for small proportions of total phenotypic variance. Thus, these results confirm the polygenic nature of body size in *P. pungitius* (Laine *et al*., 2013). However, our analysis of pelvic spine length and body depth revealed a different picture: we found little phenotypic and additive genetic variance underlying these two traits despite moderate heritabilities. Regressive evolution of the pelvic apparatus (hereafter *pelvic reduction*) has been much studied in stickleback fishes (Bell *et al.,*, 1993; Gibson, 2005; Chan *et al*., 2010; Xie *et al.,* 2019; Kemppainen *et al.,* 2021). Based on the results of Kemppainen *et al*. (2021) and the current study, it is clear that the pelvic reduction is a heterogeneous process in *P. pungitius,* and that the heritability of this trait varies from one population to another. Therefore, it is possible that the alleles (or combination of alleles) underlying pelvic variation are not segregating in our Helsinki population, in turn explaining the low additive variance of this trait. Alternatively, it is possible that the pelvic phenotype in this population is close to an adaptive optimum where stabilizing selection has depleted most of the additive genetic variance underlying this trait (Fisher, 1958; Karavolias *et al*., 2020). Whether body depth is under a similar selective scenario in *P. pungitius* is however unknown, and additional studies of selection intensity and in-depth analyses of the genetic architecture of these two traits (*e.g.*, with genome wide association study) may shed more light on their genetic bases in the Baltic Sea population of *P. pungitius*.

Interestingly, our chromosome partitioning analyses revealed heterogeneous chromosome contributions to the overall phenotypic variance for all three morphological traits, between all crosses. This result was further supported by different QTL underlying phenotypic variation between crosses, for all three traits. Differences in the genetic architecture of the same trait between these crosses indicate that different alleles govern phenotypic variation in the three different pond populations despite their adaptation to the same environment. As recently demonstrated by Kemppainen *et al*. (2021) and Fang *et al*. (2021), marine populations of *P. pungitius* display relatively high level of genetic isolation-by-distance resulting in substantial population structure in the sea. Consequently, and conversely to the classical model of parallel evolution described in the three-spined stickleback (*Gasterosteus aculeatus*; Colosimo *et al*., 2005), repeated colonization of similar freshwater habitats by *P. pungitius* was likely initiated by populations with different allele frequencies at causal loci underlying traits under selection. The likelihood of parallel evolution is therefore expected to be low among freshwater populations of *P. pungitius*, as was demonstrated for pelvic phenotypes by Kemppainen *et al.,* (2021) and further suggested by the current study along with evidence for genetic non-parallelism in the two additional morphological traits. This important result further reinforces the role of non-parallelism in polygenic adaptation among populations evolving in similar habitats (Barghi *et al*. 2020).

### Possible caveats

We found that genetic variances in our simulation study were encompassed in the HPD intervals of the animal models, but was also slightly underestimated. Although we do not have a clear explanation for the observed underestimation of *V_A_*, this result should not stem from technical issues associated with our animal models (i.e., convergence of MCMC chains, Fig. S4) and rather suggests a potential lack of statistical power to estimate *V_A_* from the proposed designs. Nevertheless, our results further highlight the importance of variance in genetic relatedness in the estimation of variance parameters.

Other sources of variance such as common environmental and parental effects may increase phenotypic similarity within full-sib families and attenuate between-family differences (Lynch & Walsh 1998). Although our data did not allow us to specifically account for parental effects, previous experimental findings of Ab Ghani *et al*. (2012) found that maternal effects on body size in both marine and pond populations of *P. pungitius* were negligible. Nonetheless, other type of experimental designs, such as paternal half-sib design (Lynch & Walsh 1998, Falconer & McKay 1996), able to yield estimates free from maternal effects would be preferable. As for the common environmental effects, estimation of genetic variance from within-family variation using the actual IBD relationships between full-sibs is not expected to be affected by such effects (Visscher *et al*. 2006). Although the current manuscript did not aim at comparing different methods or software capable of IBD estimation, it is worth noting that discrepancies may exist between different relatedness estimators (Ackerman *et al*. 2017, Weir & Goudet 2017) and that the estimation of quantitative genetic parameters from GRM-based models may require the use of several relatedness coefficients. Furthermore, the computing cost of Bayesian sampling associated with large matrices such as the GRMs used in the present paper could quickly become prohibitive with large sample sizes. Although providing a formal comparison of different R packages/software in handling such matrices was beyond the scope of this manuscript, we refer interested readers to other statistical tools such as *BGLR* (Perez & de los Campos, 2014) or *brms* (Bürkner 2017) and the recent work by de Villemereuil (2018) on the matter.

### Conclusions and prospects

In conclusion, we have explored the utility of a quantitative genomics approach in obtaining estimates of quantitative genetic parameters for non-model organisms and demonstrated that this approach should be undertaken carefully when sampling wild individuals, particularly when low variance in relatedness is expected in the study populations. Rather than maximizing sample size only, we suggest that such studies should consider sampling multiple local demes or multiple family groups to increase the variance in relatedness among individuals. While the applicability of this approach may be limited for organisms that are not amenable to be raised in laboratory/mesocosm conditions, it could be readily used for instance in many plant, insect, and fish systems to obtain estimates of quantitative genetic parameters from multiple populations simultaneously. However, since the accuracy of genomic heritability method depends on the variance in genome wide IBD (Xu, 2006; Visscher, 2009), the genomic characteristics of the model system will influence the utility and required sampling design in obtaining parameter estimates with sufficient accuracy and precision.

## Acknowledgements

We thank Federico Calboli and Ari Löytynoja for constructive comments on the earlier version of the manuscript. We thank Gabor Herczeg, Abigel Gonda, Yukinori Shimada, Mirva Turtiainen, Chris Eberlein, Takahito Shikano, Laura Hänninen, Kirsi Kähkönen, Miinastiina Issakainen and Sami Karja for help in fish breeding and DNA extractions, and Jing Yang for help with fish phenotyping. We thank Mikko Kivikoski for helpful discussion for estimation of IBD proportions. The computing resource support from CSC - the Finnish IT Center for Science Ltd administered by the Ministry of Education and Culture, Finland is gratefully acknowledged. Our study was supported by the Academy of Finland (grant nos. 129662, 134728 and 218343 to J.M.). Partial support was also received from the NSFC/RGC Joint Research Scheme sponsored by the Research Grants Council of the Hong Kong Special Administrative Region, China and the National Natural Science Foundation of China (Project No. N_HKU763/21).

## Ethical note

All experimental protocols were approved by permission (ESLHSTSTH223A) form the National Animal Experiment Board, Finland.

## Data availability

Raw sequence reads for the F_2_ crosses have been submitted to NCBIs short[read archives with accession nos.: PRJNA673430 and PRJNA672863. All data and code necessary to replicate the analyses presented in the manuscript will been deposited in the Dryad repository and R scripts are currently available at: https://github.com/afraimout/Relatedness/

## Author contributions

A.F, J.M. Z.L M.S. and P.R. conceived the study; P.R. contributed to pre-processing and getting the genotype data; A.F. analyzed the data; F.G. performed all computer simulations; J.M. and A.F. led the writing of the manuscript. All authors contributed critically to the drafts and gave final approval for publication.

## Notes

### Competing Interest Statement

The authors have declared no competing interest.

### Summary of Updates

This revision includes a new comprehensive simulation study.

