## Supplementary material for "Dissecting the genetic architecture of quantitative traits using genome-wide identity-by-descent sharing"

**Table of Contents:**

| **Supplementary methods** | **Page 2** |
| --- | --- |
| **Figure S1. Heatmap of genetic relatedness in the simulated**  **wild datasets.** | **Page 3** |
| **Figure S2. Distribution of IBD proportion in Helsinki crosses.** | **Page 4** |
| **Figure S3. Chromosome-wide heritability as a function of**  **chromosome length.** | **Page 5** |
| **Table S1. Summary statistics of the main quantitative genetics**  **parameters in sticklebacks** | **Page 6** |
| **Table S2. Chromosome length and theoretical**  **standard deviation of IBD sharing.** | **Page 7** |

**Supplementary methods**

*Inter-population crosses and fish husbandry*

For the Helsinki x Rytilampi cross (HEL x RYT), F_1_ parents produced seven successive clutches yielding a total of 287 F_2_-offspring in September and October 2008. This mating procedure was followed at different dates for the Helsinki x Pyöreälampi (HEL x PYO; mating: July-Sep 2012, F_2_ breeding: July 2012-Apr 2013) and the Helsinki x Bynästjärnen (HEL x BYN; mating: Nov 2013- Jan 2014, F_2_ breeding: Nov 2013-Aug 2014) crosses which respectively produced 288 and 313 F_2_-offspring. Grandparents were collected from the wild and mated artificially in the laboratory (Divino and Shultz, 2014). F_1_-offspring were reared in 1.4-l tanks in an Allentown Zebrafish Rack Systems (Aquaneering Inc., San Diego, USA). Water temperature was maintained at 17°C (±0.5°C) and partial water changes were made regularly by replacing water in the reservoir tank connected to each rack. A 14:10 Light:Dark rhythm was maintained throughout the experiments. Following absorption of the yolk sac, fry was fed daily with *Artemia* *nauplii* and subsequently with a mixture of live *Artemia* *nauplii* and frozen *Cyclops* 29-56 days post-hatching (dph), with frozen *Cyclops* and chironomid larvae at 57-84 dph, and with frozen chironomid larvae thereafter. The feedings occurred twice a day, and the food was provided *ad libitum*. Once the F_1_-offspring were sexually mature, one male and one female from each cross were randomly chosen and allowed to mate to produce F_2_ offspring. The F_2_ clutches were reared in 1.4 L tanks in an Allentown Zebrafish rack (Aquaneering Inc., San Diego, CA) until six days post-hatching (dph). After that, the fish were transferred to four Allentown Zebrafish racks for individual rearing and fed with live *Artemia* *nauplii*. Individual fish were housed in separate 1.4 L tanks and reared in the same conditions described above for their parents.

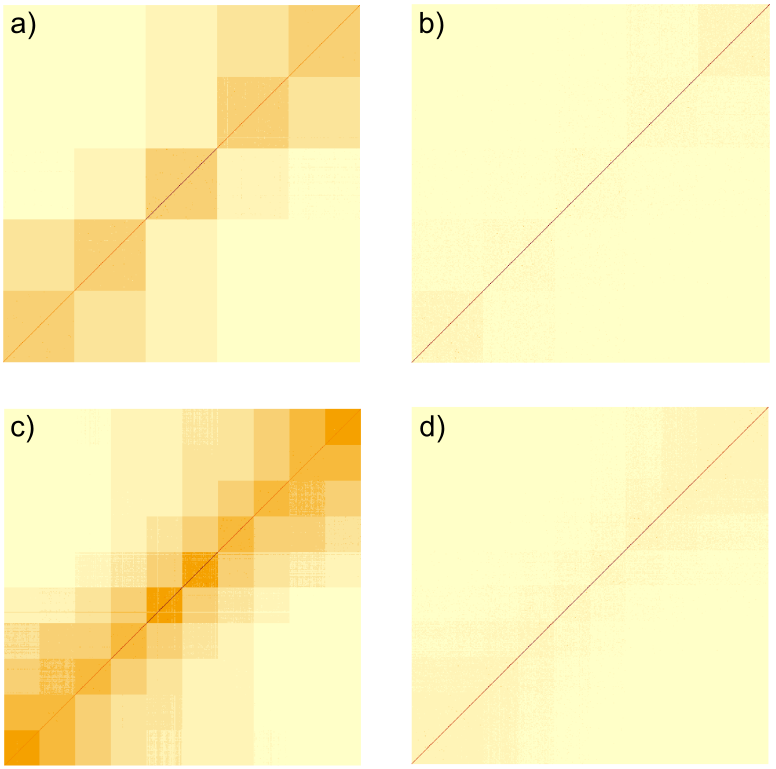

**Figure S1. Heatmap of genetic relatedness in the simulated ‘wild’ datasets.** Two meta-populations were simulated constituted by 5 (a-b) or 10 (c-d) subpopulations. Migration rate among subpopulations was set to follow four regimes from low (a, c) to high (b, d). Consequently, relatedness within subpopulations scaled negatively with migration rate and low migration populations are characterized by higher relatedness (darker color; a, c).
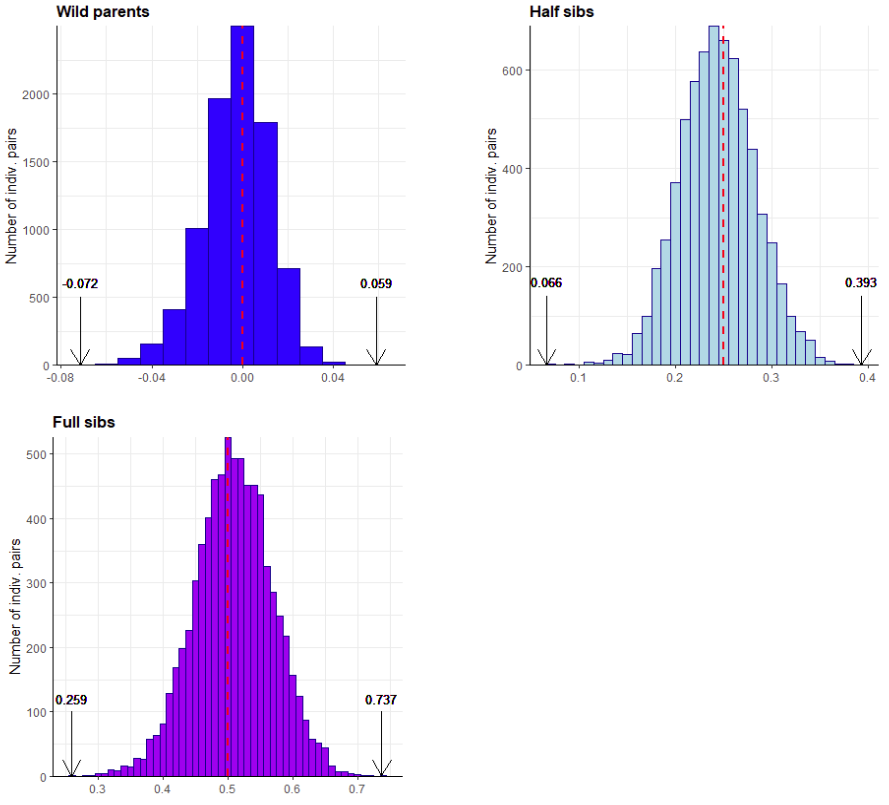

**Figure S2. Distribution of IBD proportion in Helsinki crosses.** The proportion of genome shared IBD among individuals from the Helsinki-F_1_ crosses is shown among wild parents (dark blue), full-sibs (purple) and half-sibs (light blue). For each group, the minimum and maximum values of relatedness coefficient are shown (black arrows & black text). Red vertical dashed line depicts the theoretical value of relatedness for full-sibs (0.50) and half-sibs (0.25).

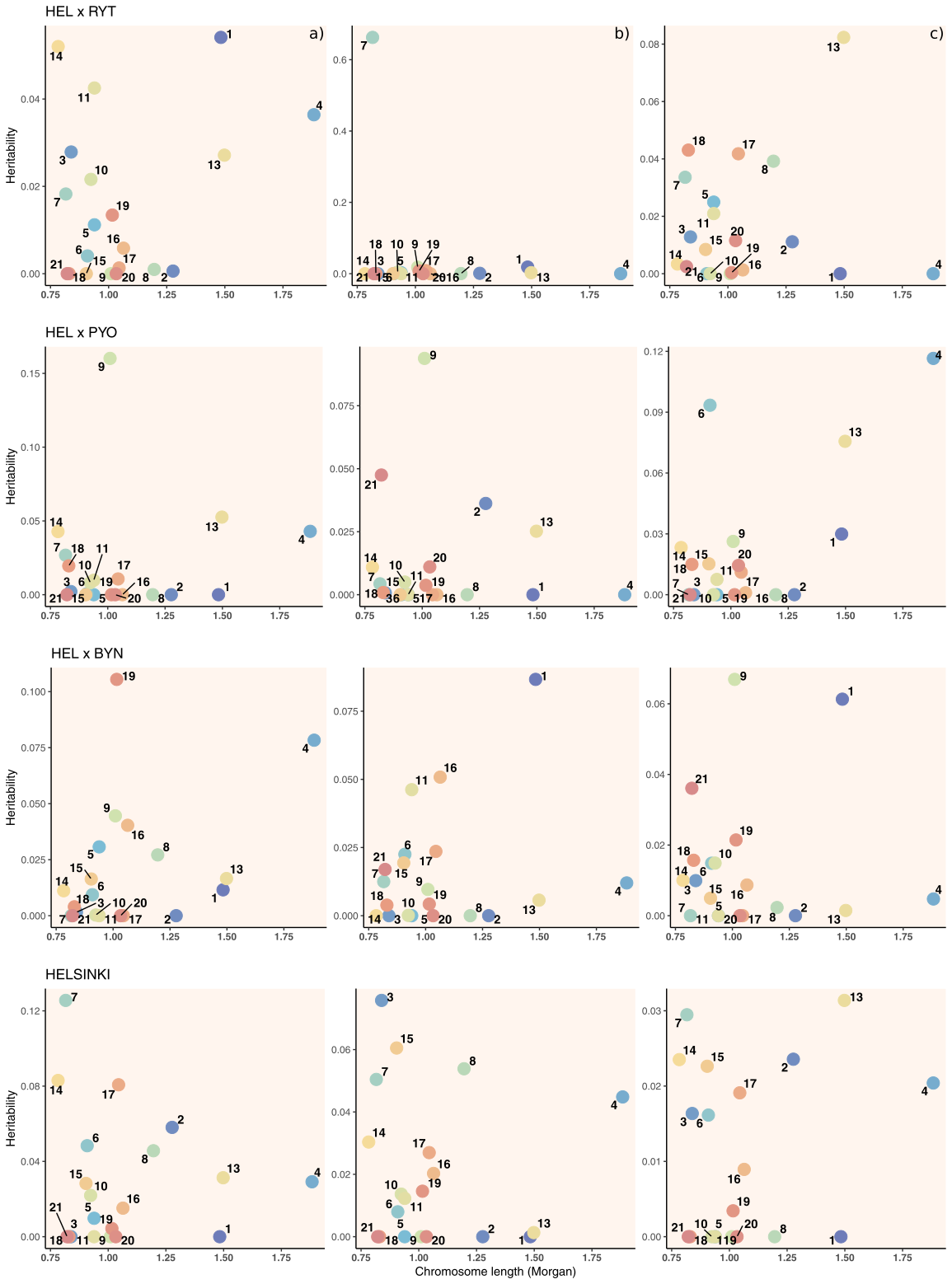

**Figure S3. Chromosome-wide heritability as a function of chromosome length.** The heritability explained by each chromosome (numbered colored points) is shown for standard length (column a), pelvic length (column b) and standard length (column c) for each cross (rows).

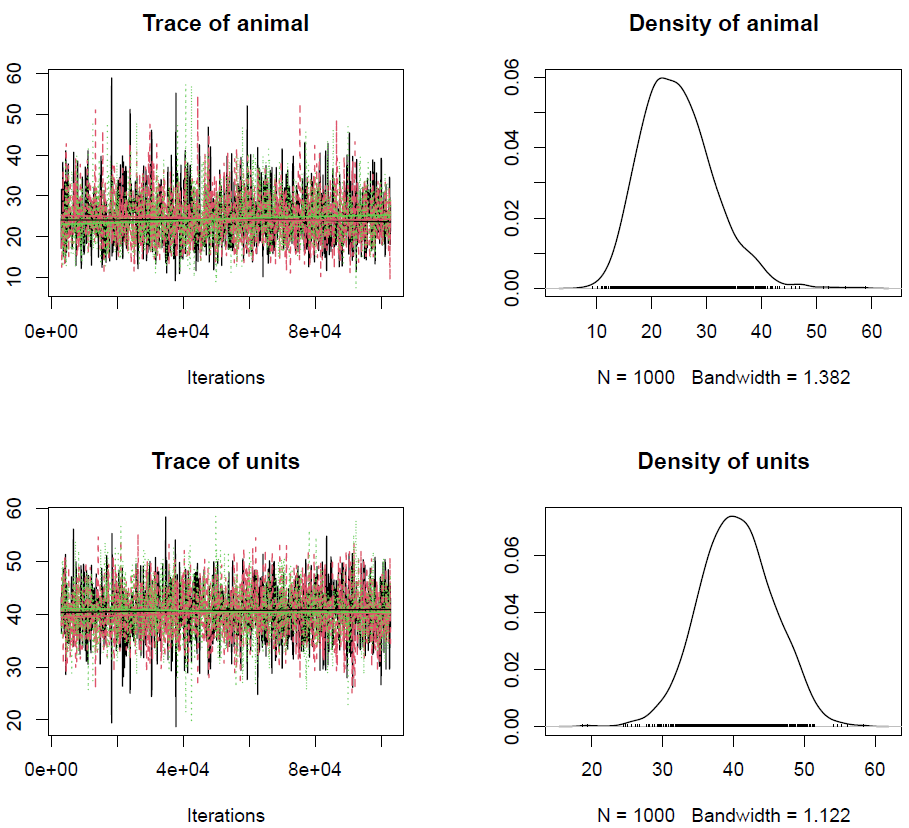

**Figure S4. Convergence of MCMC chains across models.** The convergence of MCMC chains was checked by running replicate runs of an animal model and visually inspecting the trace plots for each model (left panels, red, black and green). Overlapping of chains indicate overall convergence among MCMC runs.

**Table S1. Summary statistics of the main quantitative genetic parameters in sticklebacks.** Estimates of the total phenotypic variance (*V_P_*), additive genetic variance (*V_A_*), dominance genetic variance (*V_D_*) and heritability (*h²*) from the animal model using the PRM or GRM. Posterior median and 95% highest posterior density (HPD; in brackets) are reported for standard length (SL), body depth (BD) and pelvic length (PL) for each dataset. All variance components were corrected following Legarra (2015; see *Methods*) to provide meaningful comparisons between models and datasets.

|  |  | PRM | | | | GRM | | | |
| --- | --- | --- | --- | --- | --- | --- | --- | --- | --- |
| Cross | **Trait** | **V_P_** | **V_A_** | **V_D_** | **h^2^** | **V_P_** | **V_A_** | **V_D_** | **h²** |
| Helsinki-HS | SL | 14.999 [1.30e+01; 16.700] | 0.709 [3.77e-04; 3.744] | 8.951 [6.06e-04; 15.796] | 0.050 [2.89e-05; 0.248] | 14.995 [13.7; 16.525] | 4.099 [2.48; 6.101] | 0.632 [5.33e-04; 2.808] | 0.274 [1.64e-01; 0.380] |
|  | BD | 0.547 [4.71e-01; 0.629] | 0.104 [2.04e-04; 0.249] | 0.047 [2.78e-04; 0.290] | 0.191 [3.97e-04; 0.421] | 0.576 [5.23e-01; 0.636] | 0.173 [1.11e-01; 0.251] | 0.008 [2.67e-04; 0.061] | 0.298 [2.04e-01; 0.417] |
|  | PL | 1.765 [1.61; 1.933] | 0.021 [2.01e-04; 0.147] | 0.024 [2.68e-04; 0.245] | 0.012 [1.20e-04; 0.082] | 0.311 [2.79e-01; 0.344] | 0.087 [4.29e-02; 0.131] | 0.010 [1.40e-04; 0.055] | 0.282 [1.57e-01; 0.409] |

**Table S2. Chromosome length and theoretical standard deviation of IBD sharing.** Genome length (in both Mega base pair and Morgan scale) and theoretical expectation of the standard deviation of IBD sharing coefficient between full-sib pairs per chromosome.

| Linkage group (chromosome) | Length (Mb) | Length (Morgan) | SD (π) |
| --- | --- | --- | --- |
| 1 | 28.704 | 1.483 | 0.293 |
| 2 | 22.533 | 1.277 | 0.310 |
| 3 | 18.271 | 0.839 | 0.360 |
| 4 | 31.895 | 1.883 | 0.265 |
| 5 | 14.242 | 0.939 | 0.347 |
| 6 | 18.814 | 0.909 | 0.351 |
| 7 | 17.469 | 0.816 | 0.363 |
| 8 | 19.555 | 1.196 | 0.318 |
| 9 | 13.815 | 1.010 | 0.338 |
| 10 | 16.682 | 0.924 | 0.349 |
| 11 | 17.773 | 0.939 | 0.347 |
| 13 | 21.393 | 1.498 | 0.291 |
| 14 | 16.291 | 0.783 | 0.368 |
| 15 | 17.097 | 0.904 | 0.351 |
| 16 | 18.149 | 1.064 | 0.332 |
| 17 | 19.324 | 1.045 | 0.334 |
| 18 | 15.785 | 0.830 | 0.361 |
| 19 | 20.058 | 1.016 | 0.337 |
| 20 | 19.928 | 1.033 | 0.335 |
| 21 | 14.805 | 0.822 | 0.362 |
| genome-wide | 382.584 | 21.209 | 0.080 |
